# Comparison of Extraction Methods for the Quantification of Phytohormones from Tomato Fruits and Leaves by LC-MS/MS

**DOI:** 10.64898/2026.04.06.716604

**Authors:** Carlos A. Juarez Guzman, Linxing Yao, Corey D. Broeckling, Cristiana T. Argueso

## Abstract

Accurate, simultaneous, and efficient quantification of chemically diverse phytohormone species is a critical task towards understanding the complex system of phytohormone signaling pathways. Quantification of phytohormones with the commonly used technique liquid chromatography coupled to tandem mass spectrometry is susceptible to the influence of non-phytohormone components present in the sample, a phenomenon referred to as matrix effect. To reduce matrix effect, some phytohormone quantification methods include additional steps of cleanup of crude extracts. However, to what extent additional purification steps provide increased accuracy compared to simpler, less laborious methods is seldomly evaluated. We evaluated three previously described phytohormone extraction methods, two of which include solid-phase extraction and one that does not, in their ability to minimize matrix effect and generate accurate estimates of phytohormone species spanning six classifications, from fruit and leaf tissue of *Solanum lycopersicum* cv. Micro-Tom (tomato). Our results show that, while the methods that included solid phase extraction occasionally outperformed each other regarding matrix effect and/or recovery efficiency for broad range of phytohormones, they rarely outperformed the simpler single-phase extraction method.

**Short Abstract:** Accurate, simultaneous quantification of chemically diverse phytohormones by LC–MS/MS is frequently confounded by matrix effects, leading to the incorporation of additional purification steps. We systematically compared three published extraction protocols with or without solid-phase extraction in tomato tissues across six hormone classes. Solid-phase methods occasionally improved matrix suppression or recovery, but did not consistently outperform the single-phase approach, questioning the added value of extra cleanup steps, particularly when high-throughput is desired, as in the case of systems biology interrogations.

## INTRODUCTION

Phytohormones are small molecule signal transducers that regulate plant processes via binding to cognate receptors, inducing transcriptional and physiological reprogramming in response to environmental and developmental cues [1–3]. Many plant processes have been shown to be regulated by multiple phytohormone pathways and interactions between their components [4–8]. It is generally accepted that all plant processes are regulated not by single, insulated phytohormone pathways, but by a complex signaling system consisting of all phytohormone pathways. An enhanced understanding of phytohormone interactions is required for development of advanced crops capable of sustaining the human population in the face of drastically altered global climate patterns [9–11].

The development of *Solanum lycopersicum* (tomato) is extensively studied due to its significant agricultural importance, particularly the fruit. Much of our current understanding stems from observing changes in phytohormone concentrations across developmental stages and tissues of tomato plants [7]. Indeed, phytohormone concentration is a major determinant of signal transduction through each of the nine currently recognized *bona fide* phytohormone pathways [1]. Accurate, simultaneous, and high-throughput quantification of multiple phytohormones is a critical task for phytohormone and tomato researchers alike. This goal is especially relevant for those seeking to study the phytohormone biology through quantitative modeling. However, the chemical diversity of phytohormones, and the complex plant matrices in which they are embedded, remain major obstacles [12, 13].

Analysis of phytohormones from plant matrices is most frequently performed by ultra-performance liquid chromatography coupled to tandem mass spectrometry (UPLC-MS/MS), due to its high sensitivity and specificity, as well as its amenability to high-throughput workflows [14]. Matrix effect (ME), which is defined as the proportion of signal intensity in the presence of sample matrix compared to that in the absence of a sample matrix, can negatively impact sensitivity and specificity [15, 16]. ME arises largely through (1) incomplete isolation of target molecules from sample matrix during extraction and/or chromatographic separation and (2) enhancement or suppression of ionization of target molecules [17, 18].

Molecules contributing to ME may be removed by solid phase extraction (SPE). However, this can come at the cost of recovery efficiency (RE), which is defined as the proportion of signal intensity observed compared to an expected concentration [15, 16]. The product of ME and RE determine the overall process efficiency (PE) of a MS-based quantification workflow (i.e., the combination of extraction method and UPLC-MS/MS).

Another important consideration when developing/selecting a phytohormone quantification workflow is throughput. Detection of subtle, yet biologically meaningful quantities of phytohormones from plant tissues is dependent on complex experimental designs and statistical power. While SPE methods may result in higher PE, they are more costly and laborious, often requiring specialized equipment and more reagents than non-SPE methods, ultimately lowering throughput and overall accessibility to the phytohormone research community. A pragmatic view is that the cost incurred by SPE methods is only justified if they empirically outperform simpler, non-SPE methods in detection of multiple phytohormones and related species (i.e., precursors, catabolites, amino acid conjugates, etc.).

In this study, we evaluated the ME, RE, and PE of three previously described phytohormone extraction methods designed for use with UPLC–MS/MS, across six phytohormone classes using low-input fruit and leaf tissue from *Solanum lycopersicum* cv. Micro-Tom. Two of the examined methods utilize SPE with acetonitrile-based solutions: the method described by Cao et al. (2020) [8], which employs a Sep-Pak tC18 cartridge, and the method described by Simura et al. (2018) [9], which employs an Oasis HLB cartridge. The third method, described by Trapp et al. (2014) [10], is a relatively simpler single-phase extraction using a methanol-based solution. We conclude that the examined SPE methods rarely outperformed the single-phase method regarding ME, RE, or PE on the plant tissues tested in this study.

## EXPERIMENTAL

### Plant Materials

*Solanum lycopersicum* cv. Micro-Tom plants were grown in a climate-controlled greenhouse at the Colorado State University Plant Growth Facility, in one-gallon pots containing Promix BX soilless media. Day/nighttime temperatures ranged from 23-29/13-15 °C, respectively. Plants were watered daily with ∼100ml at 8:00am with a drip irrigation system, and fertilized as needed. At approximately 15 weeks, three fully expanded leaves and three whole fruits in the “breaker” stage were collected from three separate plants. Samples were pooled into one 15 ml conical tube according to their tissue type, comprising two final samples: a fruit sample and a leaf sample. Both samples were immediately flash frozen in liquid nitrogen and stored at −80 °C until further processing. Samples were lyophilized and pulverized to a fine powder using a ceramic mortar and pestle. Once sufficiently ground, samples were vortexed to ensure homogeneity. For non-blank samples, 2 mg (± 10%) of lyophilized tissue were used as extraction input (Fig. 1A).

**Figure 1.**
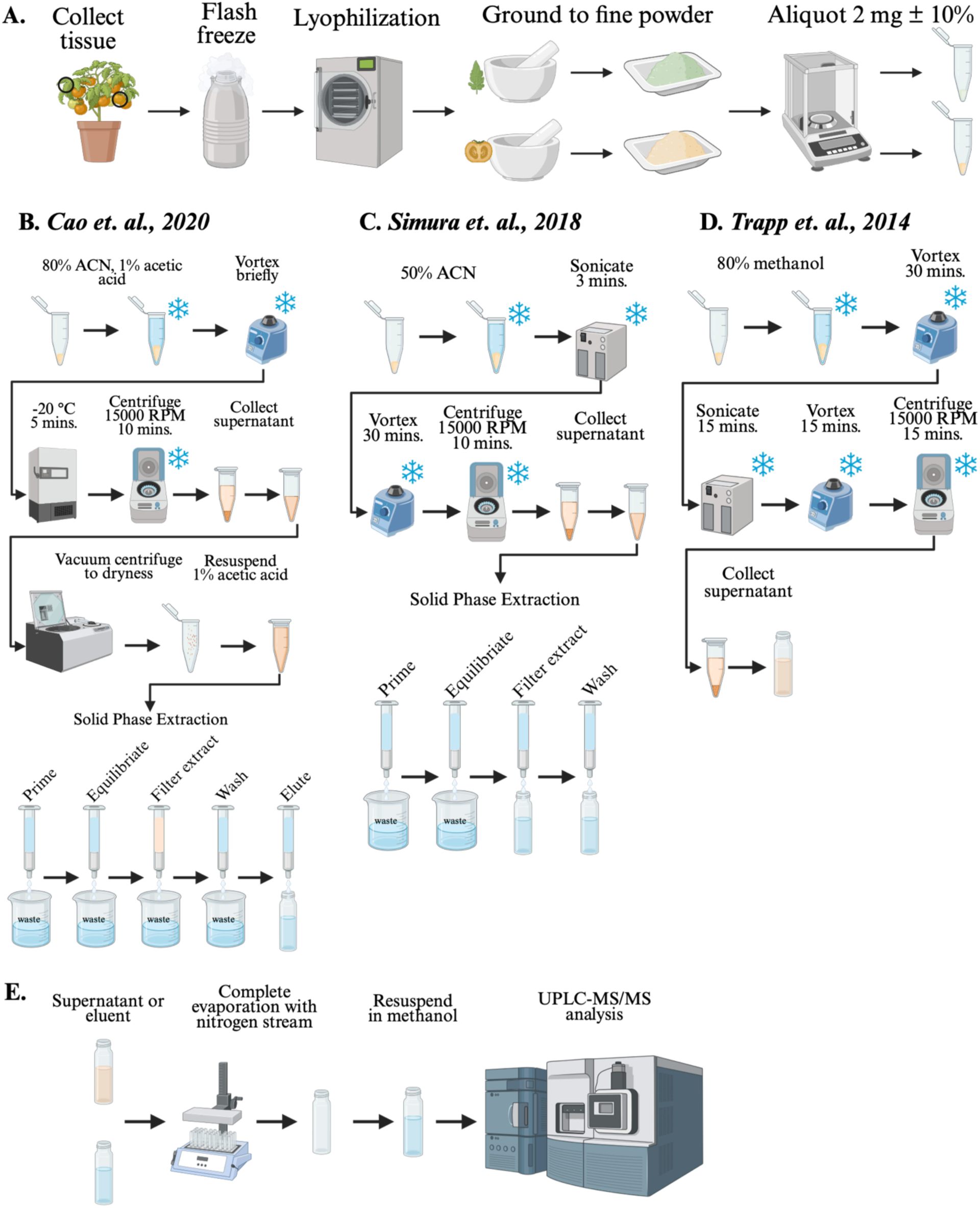
Schematic overview of sample collection (A), extraction methods (B-D), and downstream processing of extracts (D). The procedures depicted in A and E were common to all extraction methods. All extraction methods were tested in leaf and fruit tissue, although only fruit tissue is depicted in B-D.

### Reagents

All solvents used were HPLC grade and all solutions used were ice-cold at time of use. The authentic standards used in this assay including jasmonic acid (JA), salicylic acid (SA), indole 3-acetic acid (IAA), gibberellin A3 (GA_3_), gibberellin A4 (GA_4_), trans-zeatin ribose (tZR) were purchased from Sigma-Aldrich (St. Louis, MO, USA; Product numbers ADVH9B9B3801, 247588, 1.00353, 48880, G7276, and Z0375, respectively); jasmonic acid-d_5_ (JA-D_5_), salicylic acid- D_4_ (SA-D_4_), indole 3-acetic acid-D_5_ (IAA-D_5_) were purchased from CDN Isotopes (Canada; Product numbers D-6936, D-8276, and D-2203, respectively); jasmonic acid-isoleucine (JA-Ile) and trans-zeatin (tZ) were purchased from Cayman (Ann Arbor, MI; Product numbers 10740 and 13226, respectively); abscisic acid-D_6_ (ABA-D_6_), trans-zeatin-^15^N_4_ (tZ-^15^N_4_), gibberellin A_4_-D_2_ (GA_4_-D_2_), were purchased from Olchemim (Czech Republic; Product numbers 034 2721, 030 0301, and 032 2531, respectively); abscisic acid (ABA) was purchased from ChromaDex (Los Angeles, CA; Product number 032 2531).

### Phytohormone Standard Mixes

The authentic (non-labeled) standard mix contained 0.25 µg/mL of IAA, 0.1 µg/mL of tZ, 0.1 µg/mL of tZR, 25 µg/mL of ABA, 5 µg/mL of JA, 5 µg/mL of GA_3_, 10 µg/mL of GA_4_, 2.5 µg/mL of JA-Ile in 50% methanol in water. The labeled standard mix contained 0.5 µg/mL of SA-D_4_ of 10 µg/mL, 1 µg/mL of ABA-D_6_, 0.5 µg/mL of JA-D_5_, 2 µg/mL GA_4_-D_2_, 0.2 µg/mL of IAA-D_5_, 0.05 µg/mL of tZ-^15^N_4_ in 50% methanol in water. The spiking volume for either non-labeled or labeled standard mix was 10 _μ_L.

### Experimental Design

For each combination of method and tissue input, samples were spiked with known concentrations of labeled standard mix either before or after solvent extraction. All samples were spiked with non-labeled standard mix before extraction. This was done to ensure that the abundance of all phytohormones assayed could be reliably quantified across all samples. Notably, salicylic acid (SA) was not spiked into samples as we had previously observed it to be present at high abundances in fruit and leaf tissues (data not shown). Thus, the design includes three extraction method groups (Cao, Simura, and Methanol), three tissue groups (blank, fruit, and leaf), and two labeled standard spike-in groups (spiked before extraction or spiked after extraction). Five replicates for each combination of extraction method, tissue, and timing of labeled standard spike-in were included for a total of 90 samples. A schematic of this design is provided in Supplemental Figure 1.

### Cao Extractions

Cao extractions were carried out as previously described [19], with slight modifications (Fig. 1B). The extraction solvent used was aqueous 80% acetonitrile containing 1% acetic acid. Samples were vortexed vigorously and incubated at −20°C for 5 minutes. Samples were then centrifuged at 15,000 rpm for 10 minutes at 4°C. The supernatant was transferred to a new tube, vacuum centrifuged to complete dryness, and resuspended in 1 ml of a 1% acetic acid aqueous solution.

Sep-Pak tC18 (1cc) vacuum cartridges with 50 mg of sorbent were used for SPE. tC18 cartridges were primed with 1 ml of methanol and equilibrated with 1 ml of 1% acetic acid solution. Sample extracts were filtered through the cartridge, washed with 1 ml of 1% acetic acid solution, and 1 ml of 80% acetonitrile and 1% acetic acid. The eluent of 1 ml of 80% acetonitrile and 1% acetic acid was collected in a new tube and dried under a gentle nitrogen stream and resuspended in 50 _μ_L of aqueous 50% methanol (Fig. 1E).

### Simura Extractions

Simura extractions were carried out as previously described [20], with slight modifications (Fig. 1C). The extraction solvent used was aqueous 50% acetonitrile. Samples were sonicated for 3 minutes at 4°C using a Biorupter UCD-200 set to “High,” vortexed at 4°C for 30 minutes, and centrifuged at 15000 rpm for 15 minutes at 4°C. The supernatant was transferred to a new tube for further purification with an HLB cartridge.

The Oasis HLB cartridge was primed with 1 ml of methanol and 1 ml of water and equilibrated with 1 ml of 50% acetonitrile solution. Supernatant was passed through the primed HLB cartridge, and the eluent was captured in a clean tube. Additionally, 1 ml of aqueous 30% acetonitrile solution was passed through the cartridge and the eluent was collected in the same tube. The resulting 2 ml of eluent was dried under a gentle nitrogen stream and resuspended in 50 _μ_L of aqueous 50% methanol (Fig. 1E).

### Methanol Extractions

Methanol extractions were carried out as previously described [21] (Fig. 1D). The extraction solvent used was an aqueous 80% methanol solution. Samples were vortexed vigorously at 4°C for 30 minutes. Samples were then sonicated in ice cold water using a Biorupter UCD-200 set to “High” for 15 minutes after which they again were vortexed vigorously at 4°C for 30 minutes. Samples were then centrifuged at 15000 rpm for 15 minutes at 4°C. The supernatant was transferred to a new tube and dried under a gentle nitrogen stream and resuspend in 50 _μ_Lof aqueous 50% methanol (Fig. 1E).

### UPLC-MS/MS

UPLC-MS/MS analysis was performed as previously described [22] with minor modifications, on a Waters ACQUITY Classic UPLC coupled to a Waters Xevo TQ-S triple quadrupole mass spectrometer. Chromatographic separations were carried out on a Waters ACQUITY HSS T3 column (2 x 50 mm, 1.7 _μ_M). Mobile phases were (A) water with 0.1% formic acid and (B) acetonitrile with 0.1% formic acid. The LC gradient was as follows: time = 0 min, 1% B; time = 0.65 min, 1% B; time = 2.85 min, 99% B; time = 3.5 min, 99% B; time= 3.55 min, 1% B; time =5 min, 1% B. Flow rate was 0.5 mL/min and injection volume was 3 _μ_L. Samples were held at 6°C in the autosampler, and the column was operated at 45°C. Mass detector was operated in ESI+ and ESI- mode. The capillary voltage set to 0.7 kV. Inter-channel delay was set to 3 msec. Source temperature was 150°C and desolvation gas (nitrogen) temperature 450°C. Desolvation gas flow was 1000 L/h, cone gas flow was 150 L/h, and collision gas (argon) flow was 0.15 mL/min. Nebulizer pressure (nitrogen) was set to 7 Bar. The MS acquisition functions were scheduled by retention times. Autodwell feature was set for each function and dwell time was calculated by Masslynx software (Waters) to achieve 12 points-across-peak as the minimum data points per peak. The retention time, MRM transitions, cone and collision energy of each compound can be found in Table 1.

**Table 1.** MS Parameters used for detection and quantification of phytohormones.

| ESI mode | Precursor | Precursor (m/z) | Product (m/z) | Dwell (s) | Cone energy | Collision energy | RT (min) |
| --- | --- | --- | --- | --- | --- | --- | --- |
| Negative | SA | 137.1 | 65 | 0.047 | 34 | 22 | 1.94 |
| Negative | SA | 137.1 | 93 | 0.047 | 34 | 16 | 1.94 |
| Negative | SA-D4 | 141.1 | 97 | 0.047 | 30 | 15 | 1.93 |
| Negative | JA | 209.2 | 59 | 0.038 | 10 | 10 | 2.14 |
| Negative | JA-D5 | 214.1 | 62 | 0.038 | 10 | 10 | 2.13 |
| Negative | ABA | 263.2 | 153.1 | 0.047 | 32 | 10 | 1.97 |
| Negative | ABA-D6 | 269.2 | 159.1 | 0.047 | 32 | 10 | 1.96 |
| Negative | JA-Ile | 322.1 | 130.1 | 0.038 | 20 | 22 | 2.28 |
| Negative | JA-Ile | 322.1 | 172.1 | 0.038 | 20 | 18 | 2.28 |
| Negative | GA4 | 331.3 | 213.2 | 0.038 | 25 | 30 | 2.23 |
| Negative | GA4 | 331.3 | 257.2 | 0.038 | 25 | 25 | 2.23 |
| Negative | GA4-D2 | 333.2 | 215.2 | 0.038 | 25 | 30 | 2.23 |
| Negative | GA4-D2 | 333.2 | 259.2 | 0.038 | 25 | 25 | 2.23 |
| Negative | GA3 | 345.2 | 143.1 | 0.122 | 40 | 30 | 1.71 |
| Negative | GA3 | 345.3 | 239.2 | 0.122 | 40 | 18 | 1.71 |
| Positive | IAA | 176.1 | 103 | 0.052 | 20 | 30 | 1.9 |
| Positive | IAA | 176.1 | 130.1 | 0.052 | 20 | 20 | 1.9 |
| Positive | IAA-D5 | 181.1 | 134.1 | 0.052 | 20 | 14 | 1.89 |
| Positive | tZ | 220.2 | 119.1 | 0.052 | 10 | 32 | 1.36 |
| Positive | tZ | 220.2 | 136.1 | 0.052 | 10 | 18 | 1.36 |
| Positive | tZ-15N4 | 224.1 | 140.1 | 0.052 | 10 | 18 | 1.36 |
| Positive | tZ-15N4 | 224.1 | 206.1 | 0.052 | 10 | 10 | 1.36 |
| Positive | tZR | 352 | 136 | 0.052 | 30 | 28 | 1.36 |
| Positive | tZR | 352 | 220 | 0.052 | 30 | 18 | 1.45 |

### Data Processing

All raw data files were imported into the Skyline open source software package [23]. Chromatograms were manually inspected to ensure proper quantification of peak area for each phytohormone. Peak area and other relevant values were exported from Skyline as a CSV file for downstream analysis.

### Statistical Analysis

All statistical analysis was carried out in R. Two groups of linear regression models were fit to all 90 samples: One in which the response variable was the raw peak area of labeled standards and another where the response was the raw peak area of non-labeled standards. Within these two groups, one model was fit for each phytohormone species.

For models of labeled standards, tissue input (“Blank”, “Fruit”, or “Leaf”), extraction method (“Cao”, “Simura”, and “Methanol”), and timing of labeled standard spike-in (“Before Extraction” and “After Extraction”) and the three-way interaction across them were used as independent variables. For non-labeled standard models, only tissue input, extraction method, and their interaction were used as independent variables as all non-labeled standards were spiked into samples prior to extraction.

A third group of models was fit to the labeled standard-normalized peak areas of non-labeled standards in samples that were spiked with labeled standards before extraction. The phytohormones GA_3_, JA-Ile and tZR were standardized to GA_4_-D_2_, JA-D_5_, and tZ-^15^N_4_, all others were normalized to their labeled counterpart. Within this group, one model was fit to each phytohormone, with the independent variables tissue input (“Blank”, “Fruit”, or “Leaf”), extraction method (“Cao”, “Simura”, “Methanol”), and their interaction.

For all models, we determined samples to be “influential” if their Cooks Distance was > 4/n. We use the term influential throughout the text, acknowledging that some may consider these statistical outliers. We leave those points in the presentation for reasons of transparency.

## RESULTS AND DISCUSSION

### Calculation of Matrix Effect by Comparing Labeled Phytohormone Standards

Our design to quantify ME, RE, and PE was based on that described in Trufellie et. al. (2011) [16] and contained three types of samples, each defined by a combination of tissue input (leaf or fruit), extraction protocol, and timing of labeled standard spike-in (Supplemental Fig. 1). Type A samples were blank samples spiked with labeled standards after extraction. Matrix matched, or type B samples, were those with fruit or leaf input and spiked with labeled standards after extraction with one of the examined methods. Pre-spiked samples, or type C samples, were those with fruit or leaf input and spiked with the same labeled standards as in type B samples but before extraction (Supplemental Fig. 1). Labeled standards used are listed in Table 1, and their quantities described in the Experimental section.

The peak areas of labeled standards across all samples are shown in Figure 2. In general, samples spiked after extraction had greater amounts of labeled phytohormones than those spiked before. However, aside from outliers, we found little variability across blank, fruit, and leaf samples.

**Figure 2.**
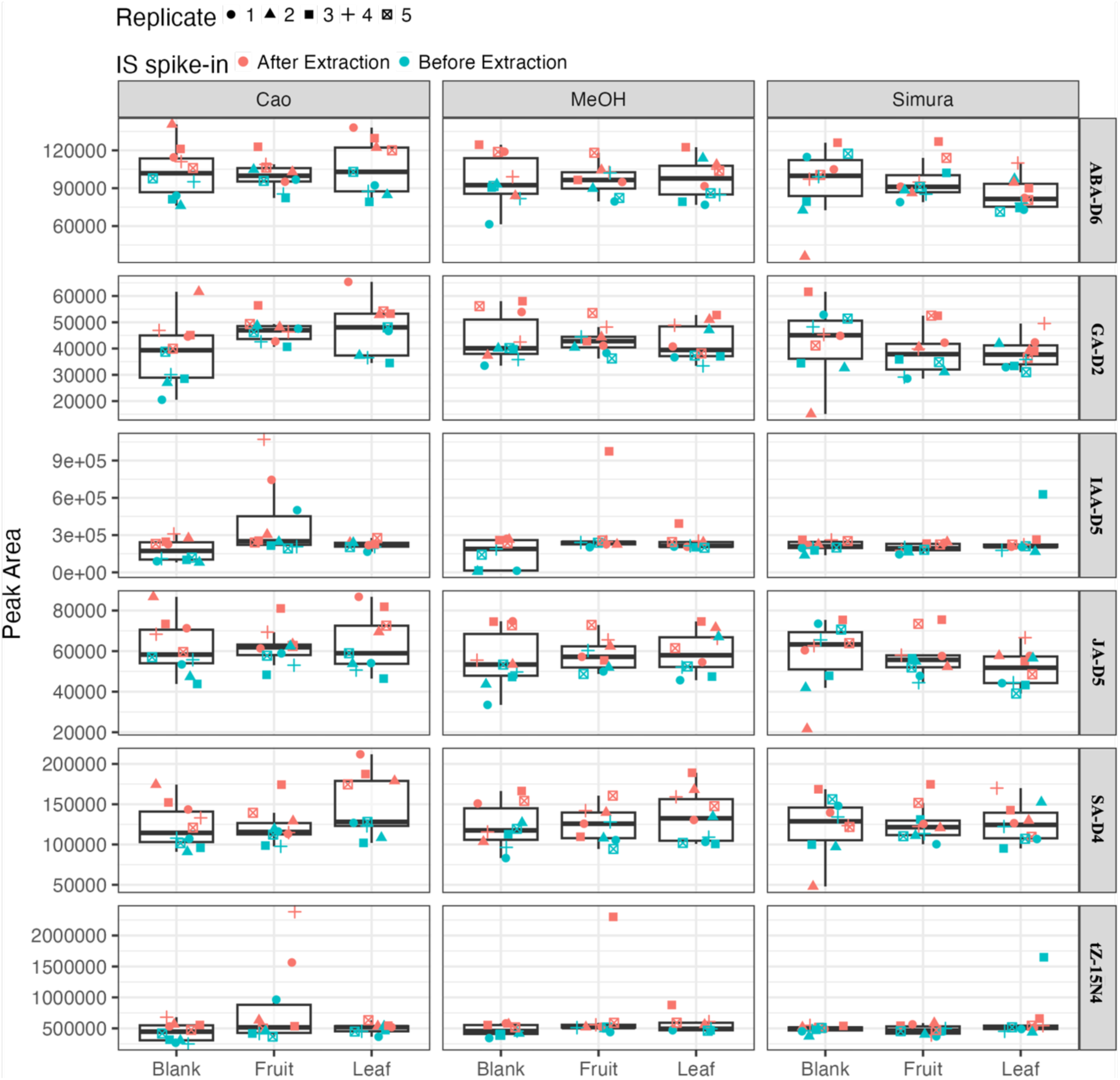
Peak area of labeled phytohormones spiked before and after extraction with different protocols tested: Each panel shows the peak area (y-axis) of labeled internal standards (IS) spiked into blank, fruit, or leaf matrices (x-axis) and extracted using one of three tested protocols: Cao, MeOH, or Simura. Colored points indicate the timing of IS addition: blue points correspond to internal standards spiked before extraction; red points correspond to standards spiked after extraction. Boxplots summarize the distribution of replicate measurements for each condition. Different symbols represent five independent technical replicates.

Using the linear regression model fit to labeled standard peak area described in the Statistical Analysis section above, we calculated ME, RE, and PE by carrying out specified contrasts on the log scale and then back transformed to the response scale, yielding a ratio. ME was calculated as the difference between type B and type A samples. Since both type A and B samples were spiked with an identical quantity of labeled standards after extraction, any difference in peak area must be caused by the presence of matrix in type B samples. Any ratio that significantly differed from 1 for ME, RE, or PE quantitatively indicated the strength and direction of effect. For example, if a comparison of type A and a group of type B (i.e., a combination of extraction and fruit or leaf tissue) samples resulted in a ratio of .8 that significantly differed from 1, then that group can be said to have a 20% reduction in signal, attributed to ME. Inversely, if the same comparison resulted in a significant ratio of 1.2, then the group can be said to have a 20% increase in signal attributed to ME. Thus, a significant ratio < 1 indicates a suppressing ME and a significant ratio > 1 indicates an enhancing ME. We expected that the simpler Methanol method would result in significant suppressing ME as it involves the least amount of purification, potentially resulting in extracts with more complex matrices than those of SPE methods.

Surprisingly, we only detected significant impact of ME with the Cao method (Fig. 3A). For fruit, use of the Cao method generated an enhancing ME of 2.03 (95% CI: 1.19 – 3.47) for IAA-D_5_ and 2.00 (95% CI: 1.13 – 3.52) for t-Z-^15^N_4_. In leaf, use of the Cao method results in a ME of 1.29 (95% CI: 1.09 – 1.55) for SA-D_4_. These results may arise from signal enhancement by the matrix, contributions from native metabolites in the matrix or other experimental artifacts. It is notable that the 95% CIs for IAA-D_5_ and tZ-^15^N_4_ were larger for all extraction-tissue type combinations in fruits, and the source of this variation is unknown. Both compounds were analyzed in positive ionization mode, however the same variation was not observed in leaf tissues, making ionization mode an unlikely source of variation in the data. These results indicate that the use of the Cao method introduces an enhancing effect on ME, specific to some phytohormones, and particularly in fruits.

**Figure 3.**
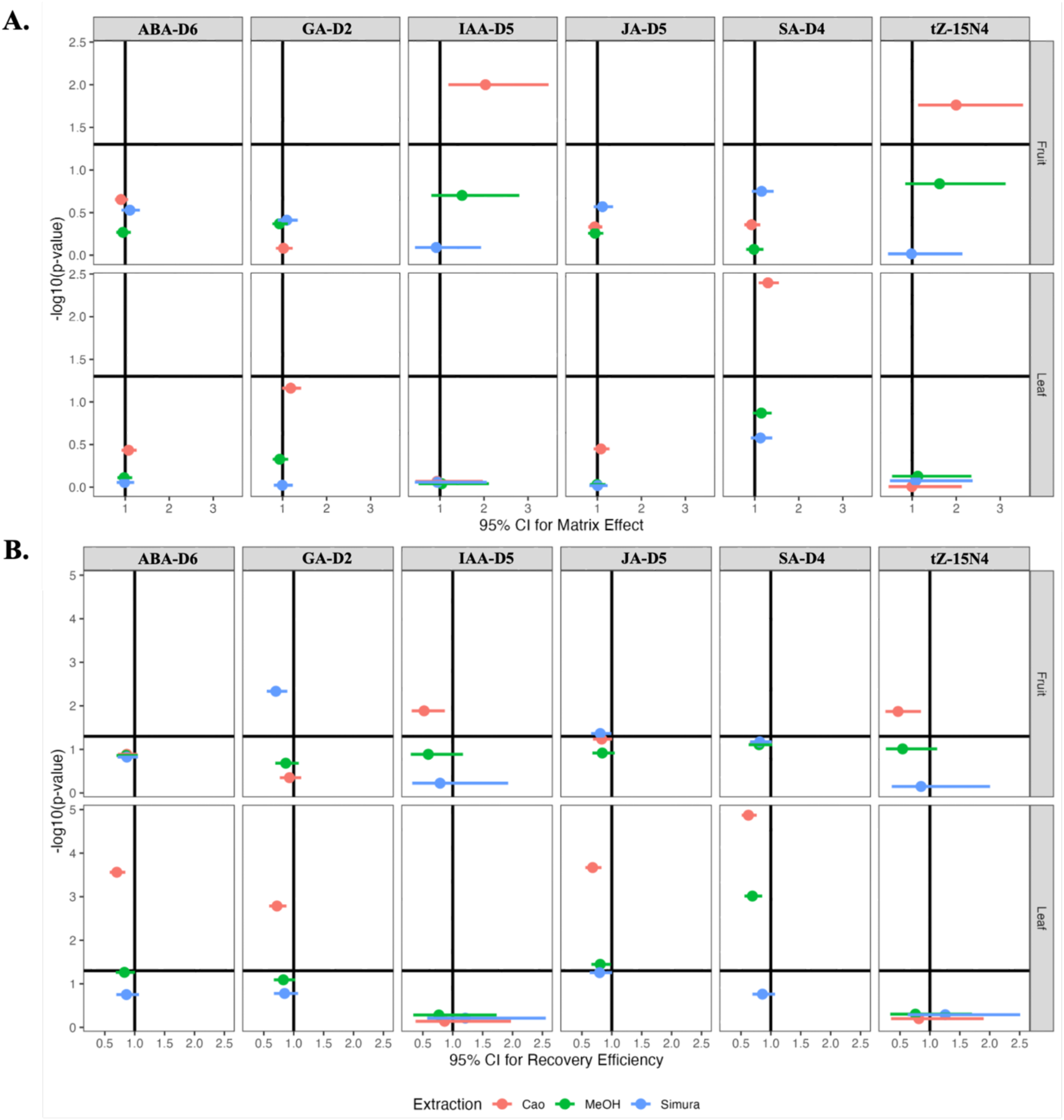
Matrix effect (ME) and recovery efficiency (RE) of labeled phytohormones: Panels summarize the estimated ME and RE, respectively, for each labeled phytohormone extracted from fruit and leaf matrices using three different protocols (Cao, MeOH, and Simura). Each point represents the mean effect size for a given protocol, with horizontal error bars indicating the 95% confidence interval (CI) of the estimated parameter. The x-axis shows the CI for ME (A) or RE (B). The y-axis displays the statistical significance as −log_10_(p-value) derived from the corresponding estimates to a value of 1. Points plotted above the horizontal reference line indicate statistically significant deviation from the null hypothesis of no matrix effect or perfect recovery.

### Calculation of Recovery Efficiency by Comparing Labeled Phytohormone Standards

RE was calculated as the difference between type B and type C samples. Both sample types contain identical tissue input and differ only in the timing of labeled standard spiking, as type B samples were spiked after extraction and type C were spiked before extraction. Thus, any difference in their peak areas for each labeled standard must be caused by loss during extraction. We expected that SPE methods would display significantly lower RE since they include an additional purification step during which phytohormones may not be fully eluted from the cartridge sorbent. This was generally true in fruit tissue, for which we detected significantly reduced RE for IAA-D_5_ and tZ-^15^N_4_ for the Cao method and GA_4_-D_2_ and JA-D_5_ for the Simura method (Fig. 3B). However, in leaf tissue, we detected significantly reduced RE for ABA-D6, GA_4_-D_2_, JA-D_5_, and SA-D_4_ for the Cao method and for JA-D_5_ and SA-D_4_ for the Methanol method.

### Calculation of Process Efficiency by Comparing Labeled Phytohormone Standards

PE was calculated as the difference between type A and type C samples on the log scale. It is possible for ME and RE to be individually statistically insignificant and yet their product, PE, to be significant. For example, the use of the Methanol protocol to extract phytohormones from fruit tissue did not lead to significant ME or RE for ABA-D_6_ or JA-D_5_, but PE was significant for both phytohormone species (< 1) (Fig. 4). Inversely, significant ME and/or RE may result in PE that does not significantly deviate from 1 by effectively canceling each other out. Several examples of this second scenario were detected. JA-D_5_ in fruit was found to have a significant RE < 1 for Simura extractions, however no significant PE was detected, suggesting that the Simura protocol generates an ME too small or too variable to be statistically detected in this study for JA-D_5_, but sufficiently large to counteract an RE of <1. The same trend was seen to a greater extent for IAA-D_5_ and tZ-^15^N_4_ in fruits extracted with the Cao protocol. Both groups had a significantly enhancing ME and significantly reduced RE yet have a PE indistinguishable from 1.

**Figure 4.**
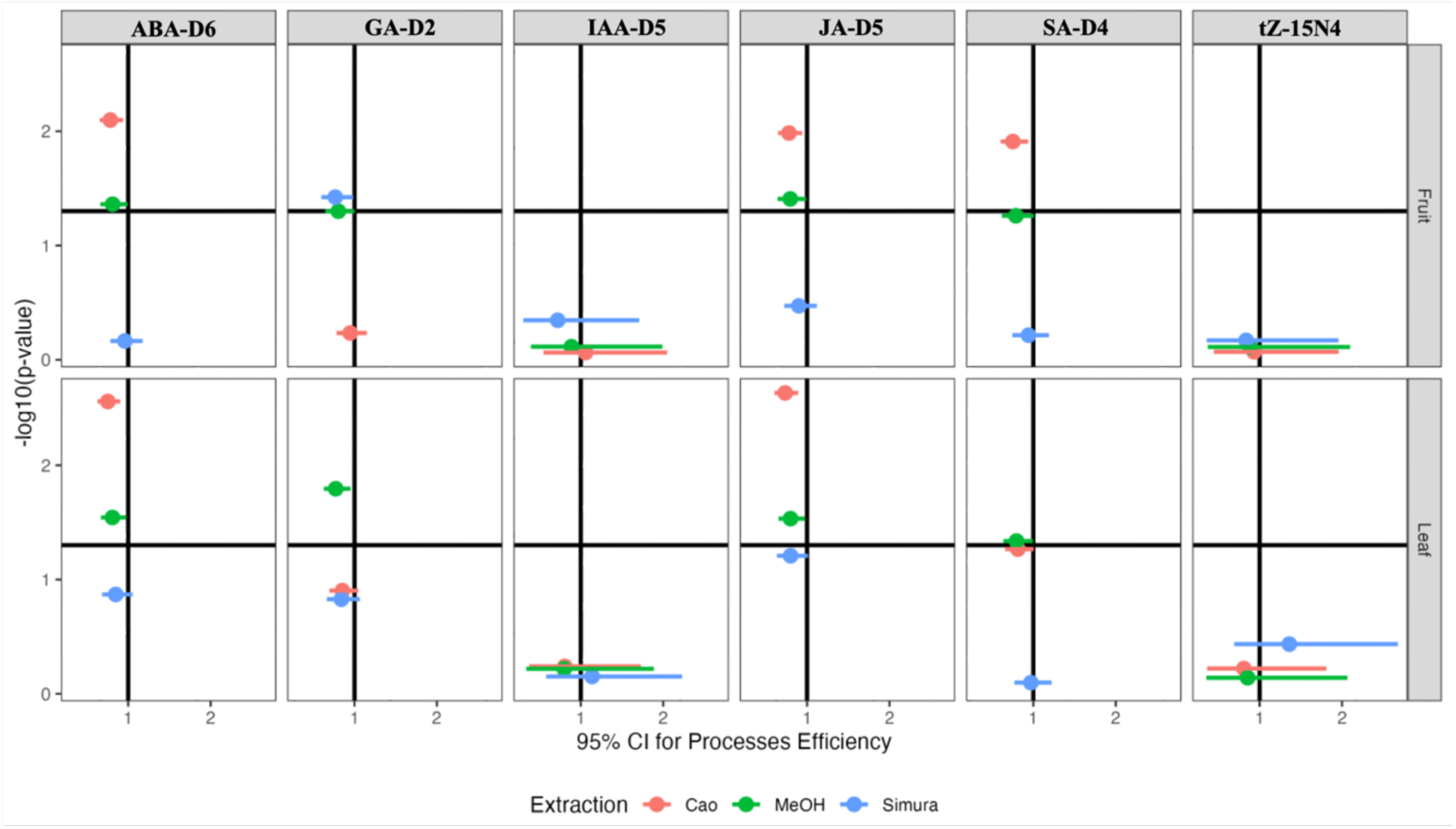
Process efficiency of labeled phytohormones: Process efficiency was calculated for each labeled phytohormone extracted from fruit and leaf matrices using the Cao, MeOH, and Simura protocols. Each point represents the estimated mean PE for a given protocol. Horizontal error bars indicate the 95% confidence interval (CI). The y-axis shows statistical significance expressed as −log_10_(p-value). Points located above the horizontal reference line indicate significant deviation from perfect PE (i.e., PE = 1). Comparisons across panels highlight the protocol and tissue-dependent variability in overall process efficiency for each phytohormone.

For labeled standards that follow the pattern described in the second scenario, it is possible that our estimates of RE are sensitive to the > 1 ME stemming from type B samples but are not present in type C samples, and thus inaccurate. This can be explained by ME that are dose dependent. However, this does not appear to be the case for the Simura method in the extraction of JA-D_5_ from fruit, for which an insignificant PE arises due to a low outlier in the type A group (Fig. 2). This outlier also inflates ME estimates for all labeled phytohormones detected via negative ESI (i.e., ABA-D_6_, GA_4_-D_2_, IAA-D_5_, JA-D_5_, and SA-D_4_) in both tissue types for the Simura method, but is unlikely to have arisen from technical error as it is not an outlier for labeled phytohormones detected via positive ESI (i.e., IAA-D_5_ and tZ-^15^N_4_), indicating that it likely arose due to injection variability.

The ME seen in IAA-D_5_ and tZ-^15^N_4_ for Cao-fruit samples may only be present at higher concentrations and/or is largely driven by outliers or statistically influential samples in the type B group. The outliers and/or statistically influential samples seen for IAA-D_5_ and tZ-^15^N_4_ are not likely due to a technical error during sample preparation, as all labeled standards were spiked together, yet they only fit this criterion for IAA-D_5_ and tZ-^15^N_4_ and are derived from the same sample injection (Fig. 2). Thus, the variation seen for IAA-D_5_ and tZ-^15^N_4_ is likely due to the unpredictable nature of ME. Positive ESI may be more susceptible to matrix enhancement and the Cao method seems to exacerbate this effect compared to the Methanol and Simura methods.

We also calculated the coefficient of variation (CV) for each extraction across fruit and leaf tissue (Table 2). CVs for IAA-D_5_ and tZ-^15^N_4_ are notably high. The CV for IAA-D_5_ is 47.38% in Cao-fruit samples and 71.06% in Simura-leaf samples. The CV for tZ-^15^N_4_ is 47.27% in Cao-fruit samples and 74.21% in Simura-leaf samples. While the mechanism underlying influential samples cannot be completely discerned, these results support that the SPE methods assessed here generate unusually high variation in PE for analytes which undergo positive ESI.

**Table 2.** Coefficient of Variation for PE: For each labeled phytohormone and tissue combination the coefficient of variation (standard deviation/mean) is presented in percentage form.

| Phytohormone | Tissue | Cao | MeOH | Simura |
| --- | --- | --- | --- | --- |
| ABA-D5 | Fruit | 9.83% | 11.44% | 9.59% |
| ABA-D5 | Leaf | 10.05% | 16.84% | 13.53% |
| GA4-D2 | Fruit | 7.54% | 8.85% | 10.35% |
| GA4-D2 | Leaf | 15.46% | 13.47% | 12.01% |
| IAA-D5 | Fruit | 47.38% | 7.81% | 12.40% |
| IAA-D5 | Leaf | 12.94% | 3.21% | 71.06% |
| JA-D5 | Fruit | 9.82% | 9.91% | 9.81% |
| JA-D5 | Leaf | 8.77% | 15.87% | 14.35% |
| SA-D4 | Fruit | 9.25% | 12.85% | 9.87% |
| SA-D4 | Leaf | 10% | 12.90% | 19% |
| tZ-15N4 | Fruit | 47.27% | 7.08% | 11.34% |
| tZ-15N4 | Leaf | 13.34% | 2.91% | 74.21% |

### Recovery of Non-labeled Standard Signal - Positive ESI: IAA, tZ, and tZR

Overall, the Simura method performed as well or better than the Cao and Methanol methods in PE for all labeled phytohormones assessed across both tissue types, except for GA_4_-D_2_ in fruit tissue, for which the Cao method minimized PE deviation from 1. However, we hypothesized that the estimates of ME for the Simura method were slightly inflated due to a low outlier in the type A group. The Cao and Methanol methods performed similarly to each other, however, the Cao method was found to be particularly prone to generating enhancing ME for labeled phytohormones detected by positive ESI. To assess the validity of these conclusions, we compared the peak area of non-labeled phytohormones across the three extraction methods in blank, fruit, and leaf samples.

Non-labeled phytohormone standards were spiked into all 90 samples in our study, alongside the addition of extraction solvent. Differences across blank samples are an estimate of differences in RE when no ME are present. Differences across extraction methods within fruit or leaf tissue represent differences in PE (i.e., ME × RE). Thus, for a given phytohormone and tissue type, if there is no difference in ME across extraction methods, then the differences seen in fruit or tissue samples should be proportional to those of blank samples. Deviation from this expected pattern would indicate differences in ME. Further, since fruit and leaf samples contain endogenous phytohormones, if blank samples from the same extraction method were found to have a greater mean peak area, then that would be indicative of a suppressing ME.

For the labeled phytohormones analyzed using positive ESI, IAA-D_5_ and tZ-^15^N_4_, we found evidence of an enhancing ME and low RE, especially for fruit tissue samples extracted using the Cao method (Fig. 3A). Because of this, we expected that blank Cao samples would contain less signal from the corresponding non-labeled phytohormones IAA and tZ, as well as the chemically related tZR, compared to blank Methanol and Simura samples. However, because we did not observe a significant PE deviation from 1 in IAA-D_5_ or tZ-^15^N_4_ PE for any method-tissue combination (Fig. 4), we did not expect significant differences in peak area for IAA, tZ, or tZR, in fruit or leaf, as this indicated that the statistically detected ME is sufficiently buffered by a statistically undetected RE.

No significant differences in blank samples for peak area of IAA, tZ, or tZR were found (Fig. 5), suggesting that the low IAA-D_5_ or tZ-^15^N^4^ RE estimated for Cao is likely an artifact of ME in type B samples. In fruit tissue, Cao samples did contain significantly greater peak area for IAA compared to the Simura method, consistent with an enhancing ME. A similar trend is seen for tZ and tZR, although there are no significant differences in peak area of tZ. Further, the Simura group has the lowest CV in fruit samples. For leaf samples, there are no significant differences for IAA, tZ, or tZR but Cao and Simura samples have a much greater CV than the Methanol method (Table 3).

**Figure 5.**
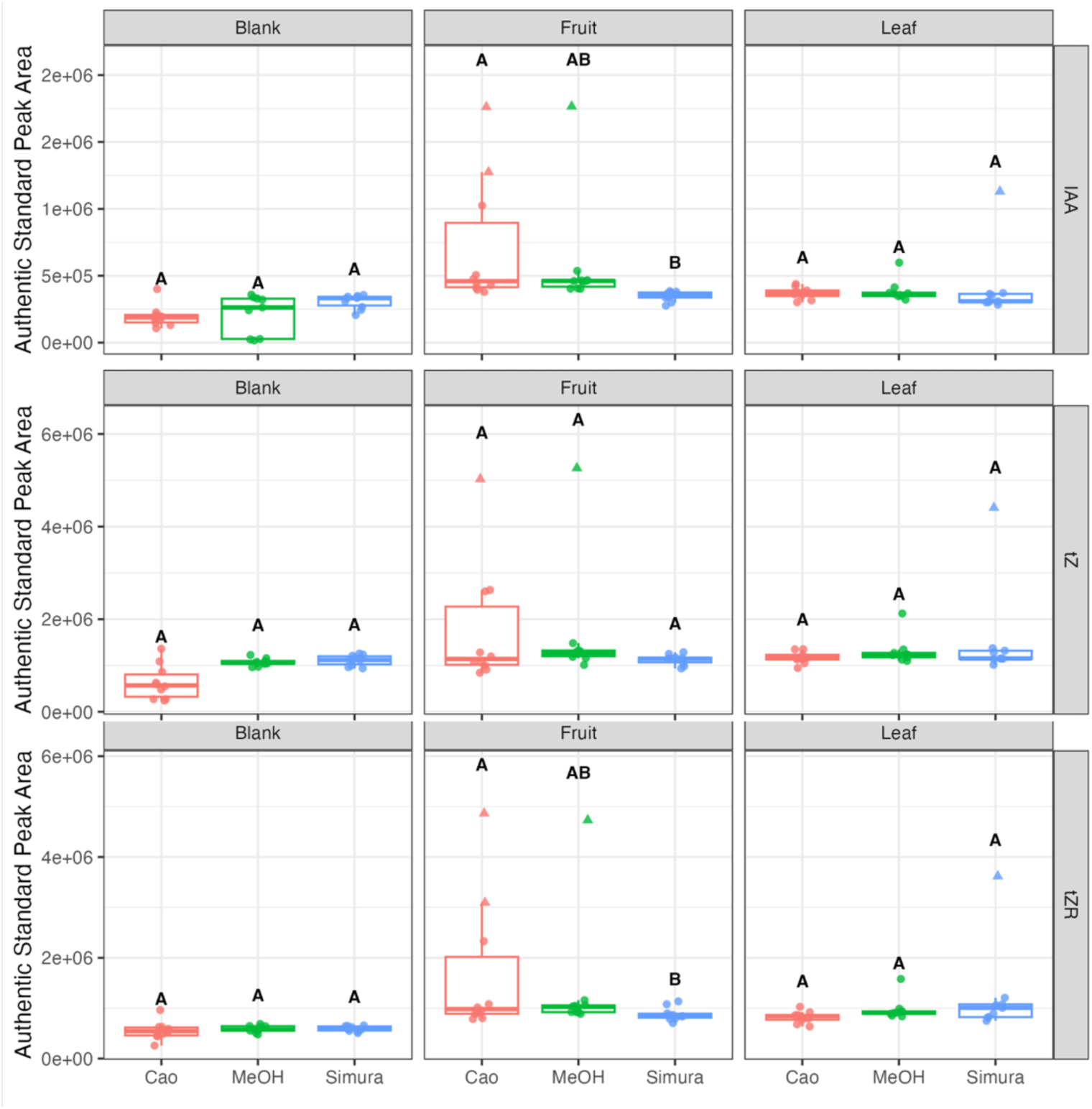
Peak Area of Phytohormones Detected with Positive Electrospray Ionization: Boxplots show the distribution of phytohormone peak areas (y-axis) for each phytohormone detected under positive ESI across extraction protocols (Cao, MeOH, and Simura) and tissue types (blank, fruit, and leaf). Letters above each box indicate results of post-hoc pairwise comparisons. Groups that do not share a letter differ significantly (P < 0.05) for the corresponding tissue–phytohormone combination. Triangles denote influential data points identified during statistical analysis (e.g., outliers or high-leverage observations).

**Table 3.** Coefficient of Variation for Non-labeled Phytohormones: For each non-labeled phytohormone and tissue combination the coefficient of variation (standard deviation/mean) is presented in percentage form.

| Phytohormone | Tissue | Cao | MeOH | Simura |
| --- | --- | --- | --- | --- |
| ABA | Fruit | 11% | 9.56% | 12.41% |
| ABA | Leaf | 54.03% | 11.62% | 13.74% |
| GA3 | Fruit | 13.65% | 11.19% | 13.09% |
| GA3 | Leaf | 48.64% | 11.59% | 14.87% |
| GA4 | Fruit | 11.59% | 9.44% | 11.70% |
| GA4 | Leaf | 38.94% | 11.57% | 13.25% |
| IAA | Fruit | 67.55% | 71.76% | 10.30% |
| IAA | Leaf | 62.41% | 21.28% | 65.37% |
| JA | Fruit | 11.14% | 10.70% | 12.48% |
| JA | Leaf | 54.90% | 13.04% | 13.30% |
| JA-Ile | Fruit | 10.19% | 10.63% | 12.26% |
| JA-Ile | Leaf | 45.55% | 12.26% | 12.63% |
| SA | Fruit | 28.25% | 28.06% | 34.85% |
| SA | Leaf | 44% | 29.69% | 42.30% |
| tZ | Fruit | 75% | 77.10% | 9.72% |
| tZ | Leaf | 64.63% | 23.69% | 69.30% |
| tZR | Fruit | 81.59% | 86.47% | 14.95% |
| tZR | Leaf | 86.46% | 23.38% | 72.07% |

The differences observed are modest, however, variation caused by ME is not typically of biological interest and can be minimized by selecting the optimal extraction method and internal standard based on the phytohormones and tissue of interest. For IAA, tZ, and tZR, the Simura extraction method minimizes ME and CV in fruit tissue, while the Methanol method minimizes ME and CV in leaf tissue.

### Recovery of Non-labeled Standard Signal - Negative ESI: ABA, GA_4_, GA_3_, JA, JA-Ile, SA

Non-labeled SA was not spiked into any samples and accordingly, the blank groups for SA show no significant differences (Fig. 6). In fruit samples, however, the Simura group was found to have significantly more SA than the Cao group, but not the Methanol group, and there was no significant difference between the Cao and Methanol groups. Further, the CV of SA in the fruit for the Simura is greater than for Cao or Methanol samples (Table 3). Together, these observations suggest that the Simura method generates a slight enhancing ME for SA in fruit. This ME is not observed for leaf tissue as all three methods generate a statistically indistinguishable SA peak area that is greater than in blank samples.

**Figure 6.**
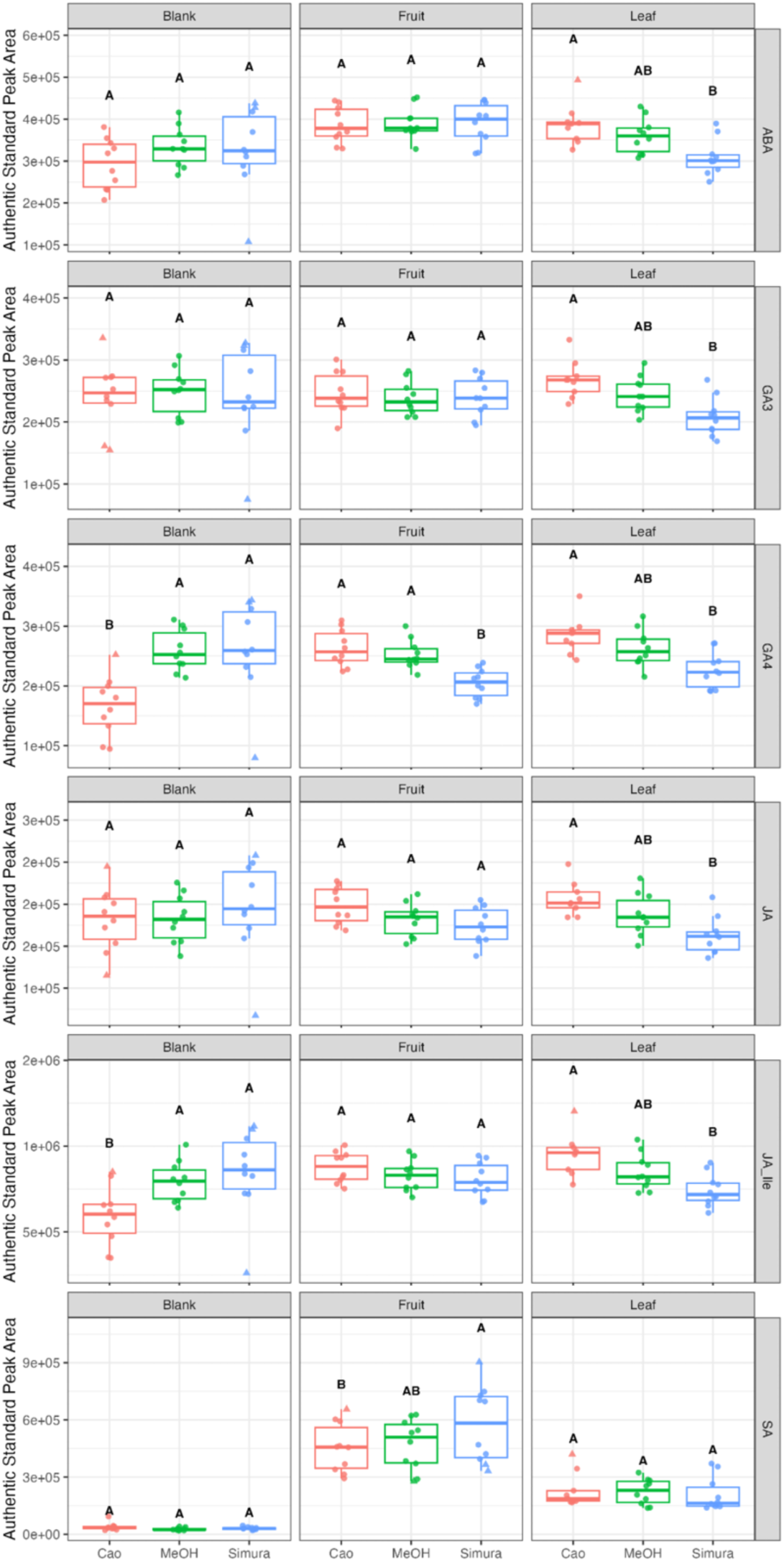
Peak Area of Phytohormones Detected with Electrospray Ionization: Boxplots display the distribution of phytohormone peak areas (y-axis) for each phytohormone quantified under negative ESI across extraction protocols (Cao, MeOH, and Simura) and tissue types (blank, fruit, and leaf). Letters above boxplots indicate results of post-hoc pairwise comparisons. Groups that do not share a letter differ significantly (P < 0.05) for the corresponding tissue–phytohormone combination. Triangles denote influential data points identified during statistical analysis (e.g., outliers or high-leverage observations).

No significant differences in peak area for ABA, GA_3_, and JA in blank samples were detected across extraction methods or fruit tissue, indicating similar PE. However, mean peak area of blank Cao samples for ABA and JA is slightly, albeit insignificantly, lower than for Methanol and Simura but not to the same extent in fruit tissue, potentially reflecting a small enhancing ME. Further, the mean peak area of JA in the Simura-fruit group is lower than that of the Simura-blank group, likely reflecting a small, suppressing ME as fruit and leaf samples contained at least as much phytohormone as the blank samples. A similar trend for the Cao and Simura samples was observed for GA_4_ and JA-Ile. The Cao group had significantly less peak area for blank samples than the Methanol or Simura group but had as much or more peak area in fruit, while the Simura group had less peak area for GA_4_ in fruit than the Cao or Methanol methods. For both GA_4_ and JA-Ile, the Simura-fruit group had a lower mean peak area than in the Simura-blank sample, although we did not assess if this difference was significant, the trend is consistent with a suppressing ME effect.

In leaf tissue, peak area of ABA, GA_3_, GA_4_, JA, and JA-Ile all displayed a similar pattern across extraction methods: The Cao group has significantly more peak area than the Simura, but does not differ from the Methanol group. This result is consistent with the trend in data from fruit samples. Generally, for phytohormones detected with negative ESI, the Cao method has reduced RE and generates an enhancing ME while the Simura method generates a suppressing ME.

We were not able to precisely quantify ME, RE, and PE for non-labeled phytohormones as we were for labeled phytohormones due to the limitations of our study design. However, differences in ME and RE across the extraction methods and phytohormones assessed here are likely to be small. This is supported by the observation that mean peak area of Methanol samples does not significantly differ from that of Cao or Simura samples for any phytohormone detected with negative or positive ESI for fruit or leaf tissues, even when the Cao and Simura groups do significantly differ from each other.

### Performance of Labeled Phytohormone Standards in Controlling Matrix Effect

A common strategy to account for difference in ME, RE, and PE across experimental groups is to spike samples with known amounts of labeled phytohormone before or alongside addition of the initial extraction solvent. Theoretically, because the endogenous phytohormone and the matching labeled phytohormone are nearly structurally identical, so too are their chemical properties, making it so that they interact with their matrix in a similar fashion. The signal intensity of the endogenous phytohormone can be normalized to that of the matching labeled species, effectively removing variation caused by ME and RE. Our data support that this is indeed an effective strategy, as we found a complete lack of statistical difference across extraction methods in peak area of phytohormone species when normalized to a matching phytohormone standard (Fig. 7: ABA, GA_4_, IAA, JA, SA, and tZ).

**Figure 7.**
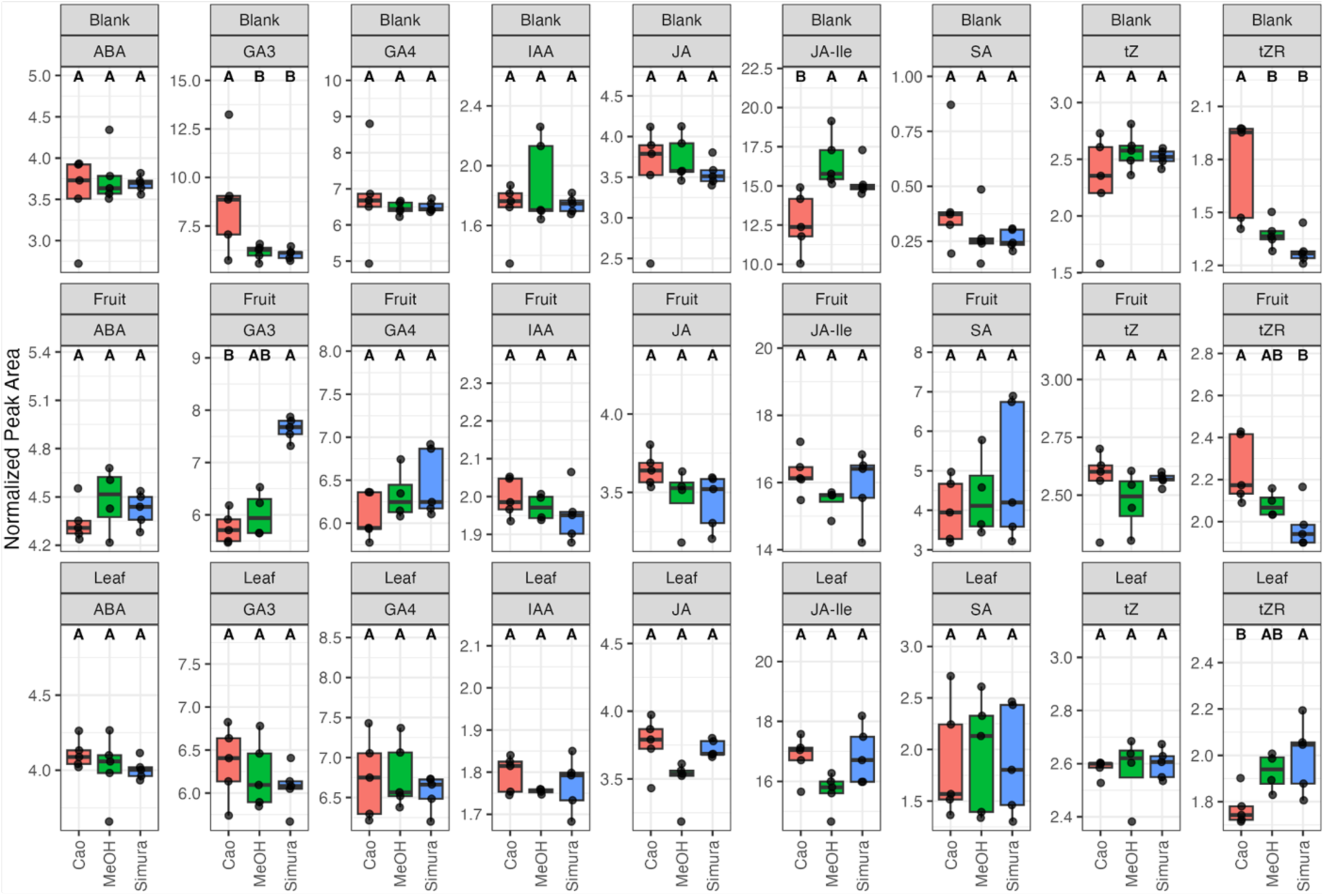
Peak Area of Phytohormones Normalized to Authentic Standards. Boxplots display the normalized peak area (y-axis) of phytohormones quantified across extraction protocols (Cao, MeOH, and Simura) and tissue types (blank, fruit, and leaf). Normalization was performed relative to corresponding labeled phytohormones. Letters above each box indicate results of post-hoc pairwise comparisons. Groups that do not share a letter differ significantly (P < 0.05) for the corresponding tissue–phytohormone combination. Black points represent individual replicates.

Many phytohormone species do not have a matching labeled phytohormone. A reasonable alternative is the use the labeled version of a closely related species, as was done here for GA_3_, JA-Ile, and tZR, which were normalized to GA_4_-D_2_, JA-D_5_, and tZ-^15^N_4_ respectively. This approach was notably less successful, as we were able to detect significant differences in peak area across extraction methods, despite normalization to a labeled, closely related phytohormone species (Fig. 7: GA_3_, JA-Ile, tZR-^15^N_4_).

### Comparative Considerations for Phytohormone Extraction Protocol Selection

The primary goal of this study was to identify the optimal extraction method for accurate and simultaneous quantification of multiple phytohormones from tomato fruit and leaf tissue. Our results indicate that the Cao and Simura methods each tend to perform better for specific phytohormones or tissue types, but neither consistently outperforms the simpler Methanol method in terms of matrix effect (ME) and recovery efficiency (RE). For accurate and simultaneous quantification of chemically diverse phytohormone species, the Methanol method appears particularly advantageous for tomato fruit and leaf. This is especially true if throughput and cost are important variables in selection of an optimal phytohormone extraction method.

While the Methanol method is suitable for broad application, the Cao and Simura methods still hold value for specialized analyses of phytohormones in tomato tissues. For example, in fruit tissue for phytohormones detected with positive ESI, CVs of Cao and Methanol methods were much greater than that of Simura samples. If a researcher was specifically interested in quantifying these molecules from tomato fruit [24, 25], then use of the Simura method would be optimal as it would reduce variation compared to the Cao or Methanol methods.

ME was estimated such that blank control samples and tissue samples were spiked with identical quantities of labeled phytohormones after extraction and the responses were compared (i.e., the post extraction method [15, 26]). Ideally, the concentrations of spiked labeled phytohormones would have mimicked endogenous concentrations of non-labeled phytohormones in fruit and leaf samples[15]. However, we found that endogenous phytohormone signal intensities from 2 +/- 0.2 mg of fruit and leaf tissue frequently approached or fell below the lower limit of the instrument’s dynamic range. This hindered reliable quantification of endogenous phytohormone concentrations and consequently, determination of optimal spiking concentrations for labeled phytohormone spike-in levels.

The studies describing the Cao and Simura methods did not assess how their respective SPE steps affect ME [19, 20]. We found little evidence for differences in ME across the three assessed extraction methods. However, this result may be an artifact of the amount of tissue (i.e., matrix) used relative to the concentrations used for the standard spike-ins. Fruit and leaf samples were expected to exhibit greater non-labeled phytohormone signal, given that the non-labeled standards are structurally identical to the endogenous phytohormones. Thus, a fruit or leaf sample would have signal from the non-labeled standard spike-ins and from the endogenously present phytohormones. An explanation for why this pattern was not always observed is that the concentrations of spiked non-labeled standards greatly exceeded endogenous levels in fruit or leaf. However, non-labeled SA was not spiked into any samples and yielded similar results to other phytohormones where Methanol performs as well as or better than Cao and/or Simura, suggesting that the spiked phytohormones did not completely obscure ME. Further, spike-in concentrations of labeled phytohormones were lower than for their non-labeled counterpart.

This study highlights the broader challenges associated with comparing extraction methods for phytohormone quantification. Published protocols typically differ in several procedural details, including solvent composition, pH, extraction volume, purification steps, and mechanical disruption [19–22, 27–30]. Because these variables interact with one another, it is difficult to determine the influence of any single factor on extraction performance. These limitations emphasize the value of community-wide efforts that promote standardized benchmarking frameworks for phytohormone extraction. Such efforts could include shared reference tissues, common spike-in levels, and consistent reporting of ME, RE, and PE. Establishing these standards would allow more direct comparisons across studies and would help identify extraction parameters that are broadly effective across plant species and phytohormone classes.

The differences in ME, RE, and PE between fruit and leaf tissue in this study demonstrate the influence of matrix composition on quantitative parameters. The metabolic diversity across plant species and tissues is substantial and includes differences in secondary metabolites, cell wall components, lipids, and pigments [31, 32]. No single extraction solvent or protocol is likely to perform equally well across all plant matrices. Expanding method comparisons to diverse tissues such as roots, flowers, developing seeds, or stress-induced organs would help reveal matrix-dependent trends that can guide solvent selection for specific biological contexts. Further extending these analyses to phylogenetically diverse species may also identify broad matrix categories or chemical features that predict extraction behavior. This information would support more deliberate and informed optimization of phytohormone quantification workflows.

Finally, these findings carry broader implications for multi-hormone and systems-level phytohormone studies. As analytical workflows increasingly aim to quantify many phytohormone classes simultaneously, generalist extraction chemistries that maintain compatibility with chemically diverse analytes become essential. Our results suggest that simple, high-throughput methods such as the Methanol extraction provide a robust foundation for such work, whereas SPE-based methods may introduce analyte-specific variability that complicates comparison across phytohormone species and different plant tissues. Continued refinement of extraction workflows, guided by evaluations across diverse plant matrices and by broader access to suitable labeled standards, will support progress in simultaneous quantification of chemically diverse phytohormone species and deepen our understanding of the phytohormonal interactions.

## Acknowledgements

We thank the Colorado State University Plant Growth Facility for their help in plant growth and care. LC-MS/MS experiments were performed in the Colorado State University Analytical Resources Core (RRID: SCR_021758).

