## Supplemental Figure 1 for "Comparison of Extraction Methods for the Quantification of Phytohormones from Tomato Fruits and Leaves by LC-MS/MS"

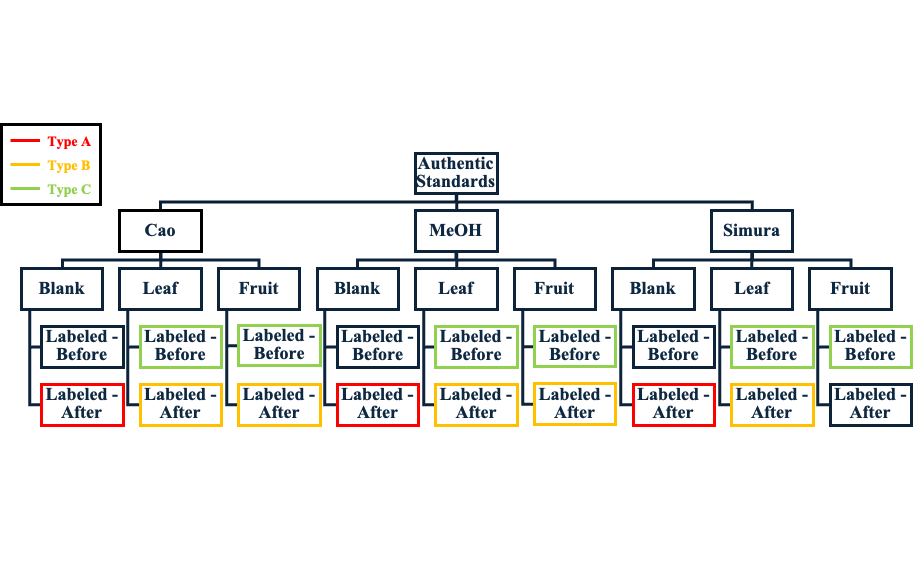

**Supplemental Figure 1.** Graph Schematic of Study Design. Type A, B, C represent the different ways that labeled standards were used.
